## Supplementary material for "A new protocol for multispecies bacterial infections in zebrafish and their monitoring through automated image analysis": S1 File Protocols

RESERVED DOI:
**10.17504/protocols.io.rm7vzjybxlx1/v1**

Désirée A. Schmitz[1,2], Tobias Wechsler[1], Hongwei Bran Li[1,3], Bjoern H. Menze[1], Rolf Kümmerli[1]

[1]Department of Quantitative Biomedicine, University of Zurich, Zurich, Switzerland;
[2]Department of Microbiology, Harvard Medical School, Boston, Massachusetts, USA;
[3]Athinoula A. Martinos Center for Biomedical Imaging, Massachusetts General Hospital, Harvard Medical School, Boston, Massachusetts, USA

Desiree Schmitz
Harvard Medical School

**Collection Info:** Désirée A. Schmitz, Tobias Wechsler, Hongwei Bran Li, Bjoern H. Menze, Rolf Kümmerli . A new protocol for multispecies bacterial infections in zebrafish and their monitoring through automated image analysis. **protocols.io**
**https://protocols.io/view/a-new-protocol-for-multispecies-bacterial-infectio-dc462yze**

**Created:** April 16, 2024

**Last Modified:** May 01, 2024

**Collection Integer ID:** 99198

**Funders Acknowledgement:**
**Swiss National Science Foundation**
**Grant ID: 310030_212266**
**Swiss National Science Foundation**
**Grant ID: 31003A_182499**

#### Abstract

The zebrafish *Danio rerio* has become a popular model host to explore disease pathology caused by infectious agents. A main advantage is its transparency at an early age, which enables live imaging of infection dynamics. While multispecies infections are common in patients, the zebrafish model is rarely used to study them, although the model would be ideal for investigating pathogen-pathogen and pathogenhost interactions. This may be due to the absence of an established multispecies infection protocol for a defined organ and the lack of suitable image analysis pipelines for automated image processing. To address these issues, we developed a protocol for establishing and tracking single and multispecies bacterial infections in the inner ear structure (otic vesicle) of the zebrafish by imaging. Subsequently, we generated an image analysis pipeline that involved deep learning for the automated segmentation of the otic vesicle, and scripts for quantifying pathogen frequencies through fluorescence intensity measures. We used *Pseudomonas aeruginosa, Acinetobacter baumannii, and Klebsiella pneumoniae*, three of the difficult-to-treat ESKAPE pathogens, to show that our infection protocol and image analysis pipeline work both for single pathogens and pairwise pathogen combinations. Thus, our protocols provide a comprehensive toolbox for studying single and multispecies infections in real-time in zebrafish.

#### Files

SEARCH

##### Protocol

NAME

Protocol (A): Zebrafish infections into the otic vesicle (2 dpf)

VERSION DC452YY6

CREATED BY

Desi
Desiree Schmitz
Harvard Medical School

OPEN →

##### Protocol

NAME

Protocol (B): Zebrafish embedding and imaging (3 dpf)

VERSION DC472YZN

CREATED BY

Desi
Desiree Schmitz
Harvard Medical School

OPEN →

##### Protocol

NAME

Protocol (C): Automated segmentation of the otic vesicle and image analysis

VERSION DC482YZW

CREATED BY

Desi
Desiree Schmitz
Harvard Medical School

OPEN →

### Protocol (A): Zebrafish infections into the otic vesicle (2 dpf)

In 1 collection

RESERVED DOI:
**10.17504/protocols.io.j8nlk8kwwl5r/v1**

Désirée A. Schmitz[1,2], Tobias Wechsler[1], Hongwei Bran Li[1,3], Bjoern H. Menze[1], Rolf Kümmerli[1]

[1]Department of Quantitative Biomedicine, University of Zurich, Zurich, Switzerland;
[2]Department of Microbiology, Harvard Medical School, Boston, Massachusetts, USA;
[3]Athinoula A. Martinos Center for Biomedical Imaging, Massachusetts General Hospital, Harvard Medical School, Boston, Massachusetts, USA

Desiree Schmitz
Harvard Medical School

**Protocol Info:** Désirée A. Schmitz, Tobias Wechsler, Hongwei Bran Li, Bjoern H. Menze, Rolf Kümmerli . Protocol (A): Zebrafish infections into the otic vesicle (2 dpf). **protocols.io** **https://protocols.io/view/protocol-a-zebrafish-infections-into-the-otic-vesi-dc452yy6**

**Created:** April 16, 2024

**Last Modified:** May 01, 2024

**Protocol Integer ID:** 99197

#### Abstract

This protocol details the zebrafish infections into the otic vesicle.

### Guidelines

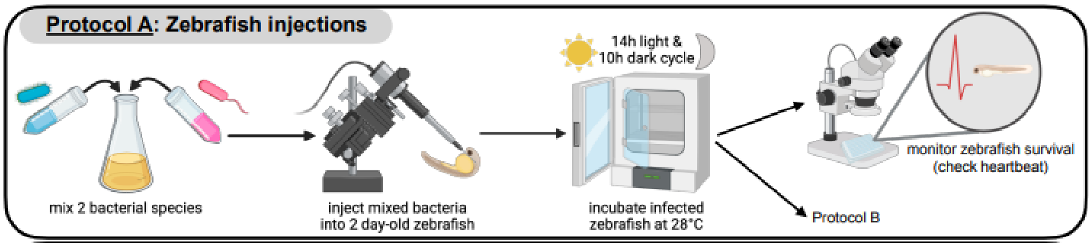

**Table SP1.** Media preparation of E3 zebrafish water (60x stock and 1x), pronase for dechorionation, PTU to avoid pigmentation, tricaine for anesthetizing.

| A | B | C |
|---|---|---|
| Solution | Amount | Chemical |
| E3 60x stock | | |
| | 34.4 g | NaCl |
| | 1.52 g | KCl |
| | 2.9 g | CaCl * 2H20 |
| | 9.8 g | MgSO4 * 2H2O |
| | | Fill up to 2000 mL with milliQ water |
| | | Stir and autoclave (1 L suffices for 60 L 1x E3) |
| E3 1x | | |
| | 165 mL | 60x E3 |
| | 10 L | milliQ water |
| Pronase | | |
| Aqueous solution stable for 1 y at -20°C | 5 g | Pronase (from Roche) |
| Shelf life: as indicated | 167 mL | milliQ water |
| 1. Mix 5 g Pronase with 167 mL with stirrer **under hood** until pronase is dissolved. 2. Store at -20°C in 1 mL aliqouts. | | |
| PTU | | N-Phenylthiourea (e.g. P7629-10 g) |
| Store powder in toxic material area (Shelf life: indefinite) | 304 mg | PTU |
| Option1: **10x** | | #ERROR |

| A | B | C |
|---|---|---|
| | | Stir overnight in fumehood (toxic!), cover in aluminum foil |
| Option2: **2500x** first dissolve in DMSO & store aliquots at 4°C | 750 mg | PTU |
| | 10 mL DMSO | Stir overnight, cover in aluminum foil |
| | | =2’500X STOCK → ADD 400 ML TO 1 L E3 (0.003% PTU) |
| **Tricaine/Mesab** | | |
| (= 4000 mg/L) | 4g | Tricaine/Mesab |
| | | Fill up with milliQ water to 1000 mL |
| | | Adjust pH at 7.0 |
| | | Store at -20°C |

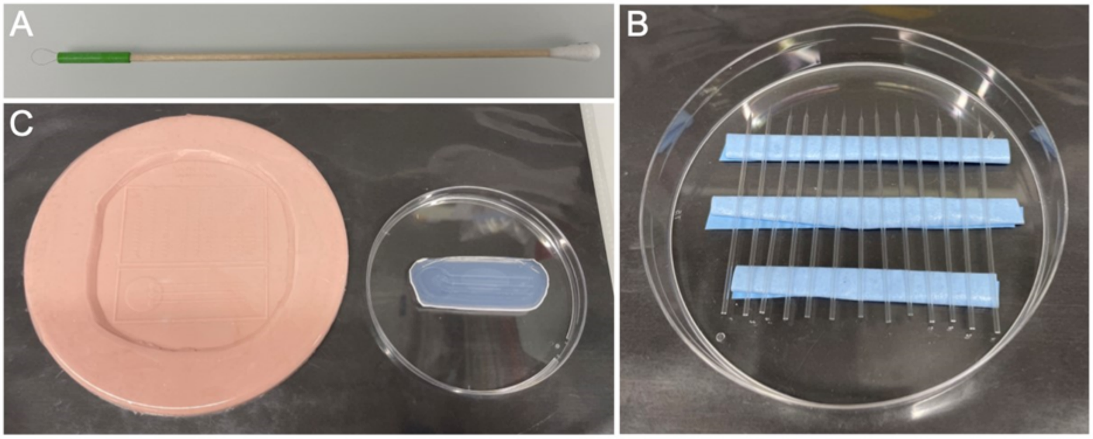

**Figure SP1**. **Material that needs to be prepared prior to injections.**
(A) Hair loop manipulator. (B) Needles. (C) Rubber mold (left) and agarose mold made with it (right).

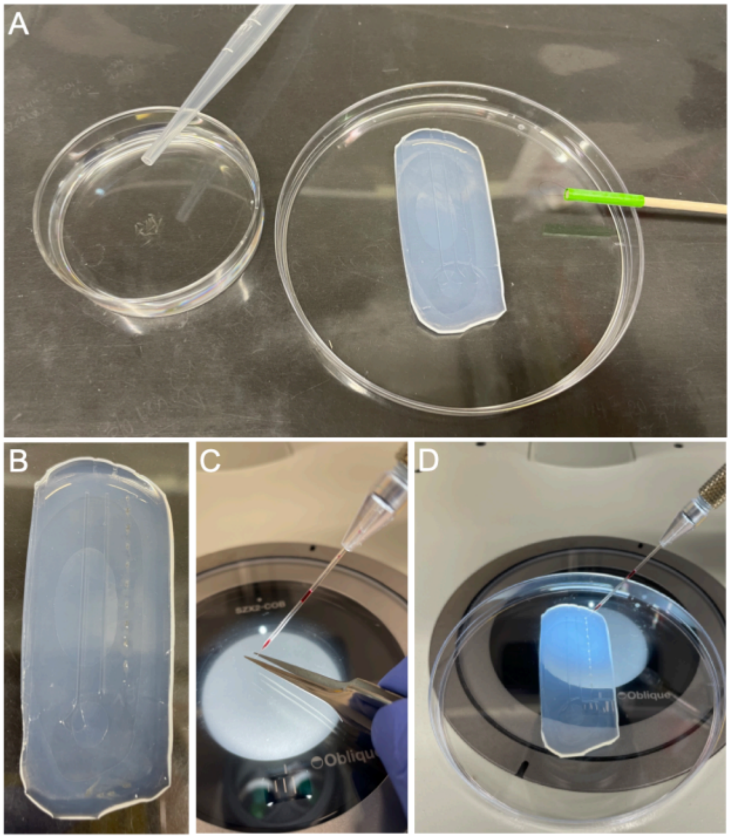

**Figure SP2. Steps illustrating the injection protocol.** (A) On the left are 2-days-post-fertilization zebrafish in a small petri dish containing an anesthetizing solution (tricaine). A Pasteur pipette can be seen on top for moving the zebrafish from the petri dish onto the agarose mold. On the right is an agarose mold (placed on an inverted lid of a petri dish) and a hair loop manipulator. (B) An agarose mold with zebrafish aligned in the rightmost channel.(C) Breaking off the needle with fine tweezers using a stereomicroscope. The needle is held by a micromanipulator. (D) Injecting zebrafish using a stereomicroscope and a micromanipulator holding a glass needle.

#### Materials

- All media were purchased from Sigma Aldrich, Switzerland unless indicated otherwise.
- All experiments in this study were conducted using lysogeny broth (LB).

**Equipment:**

For this protocol, you will need the following equipment:

- An incubator set to 37°C at 170 rpm with aeration (e.g., Infors, Multitron Standard Shaker)
- A spectrophotometer to measure optical density at a wavelength of 600 nm
- A centrifuge similar to MPW Med. Instruments, MPW-352R
- A stereomicroscope with a reticle and ruler plate in one ocular, e.g. Olympus SZX7 (final magnification of 56X)
- Micropipette puller device, for example, Sutter Instrument Co., Model Flaming/Brown P-87 with the following settings: air pressure=500, heat=609, pull=200, velocity=100, time=50
- A micromanipulator, e.g. Märzhäuser Wetzlar, MM33
- An injection pump similar to the Eppendorf FemtoJet 4i
- A temperature- and light-controlled incubator set to 28°C and a 14h light- and 10h dark-cycle, e.g., Memmert IPP110ecoPlus
- A negative rubber mold of a microstructured surface array from Ellet & Irimia (Ellett F, Irimia D. Microstructured Surface Arrays for Injection of Zebrafish Larvae. Zebrafish 2017;14:140–5. https://doi.org/10.1089/zeb.2016.1402). Their device is available for purchase at BioMEMS Core at the Massachusetts General Hospital (https://researchcores.partners.org/biomem/about).

**Table SP1.** Media preparation of E3 zebrafish water (60x stock and 1x), pronase for dechorionation, PTU to avoid pigmentation, tricaine for anesthetizing.

| A | B | C |
|---|---|---|
| Solution | Amount | Chemical |
| **E3 60x stock** | | |
| | 34.4 g | NaCl |
| | 1.52 g | KCl |
| | 2.9 g | CaCl * 2H20 |
| | 9.8 g | MgSO4 * 2H2O |
| | | Fill up to 2000 mL with milliQ water |
| | | Stir and autoclave (1 L suffices for 60 L 1x E3) |
| **E3 1x** | | |
| | 165 mL | 60x E3 |
| | 10 L | milliQ water |

| A | B | C |
|---|---|---|
| **Pronase** | | |
| Aqueous solution stable for 1 y at -20°C | 5 g | Pronase (from Roche) |
| Shelf life: as indicated | 167 mL | milliQ water |
| 1. Mix 5 g Pronase with 167 mL with stirrer **under hood** until pronase is dissolved. 2. Store at -20°C in 1 mL aliqouts. | | |
| **PTU** | | N-Phenylthiourea (e.g. P7629-10 g) |
| Store powder in toxic material area (Shelf life: indefinite) | 304 mg | PTU |
| Option1: **10x** | | =10x stock → Fill up to 1000 mL with milliQ water |
| | | Stir overnight in fumehood (toxic!), cover in aluminumfoil |
| Option2: **2500x** first dissolve in DMSO & store aliquots at 4°C | 750 mg | PTU |
| | 10 mL DMSO | Stir overnight, cover in aluminumfoil |
| | | =2’500X stock → add 400 µL to 1 L E3 (0.003% PTU) |
| **Tricaine/Mesab** | | |
| (= 4000 mg/L) | 4g | Tricaine/Mesab |
| | | Fill up with milliQ water to 1000 mL |
| | | Adjust pH at 7.0 |
| | | Store at -20°C |

#### Part 0: Material preparation

1 Media: prepare all required media according to Table SP1 (E3 zebrafish water, PTU to avoid pigmentation, pronase for dechorionation, tricaine for anesthetizing).

2 Hair loop manipulators (Fig. SP1A): tape (e.g. using 19 mm wide tape) a piece of hair as a 4-7 mm loop as an extension onto e.g. the wooden end of a cotton swab (ROTILAB, 150mm long) (see image of hairloop tool).

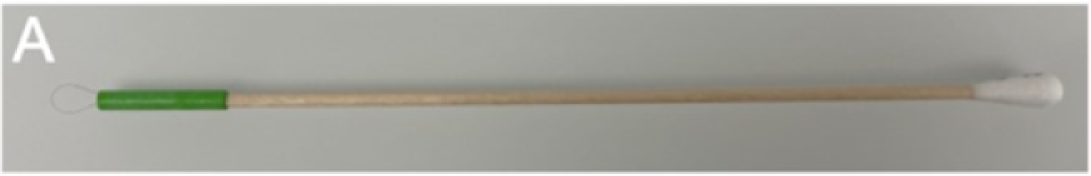

Hair loop manipulators (Fig. SP1A)

3 Needles (Fig. SP1B): Using the micropipette puller device, pull filamented borosilicate glass microcapillary injection needles (Science Products, 0.58mm diameter, GB100F-10) with the following settings: air pressure=500, heat=609, pull=200, velocity=100, time=50.

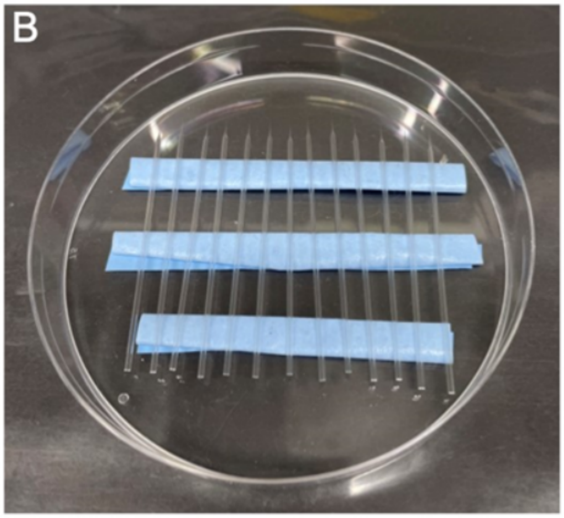

Pulled needles. (Fig. SP1B)

4 Agarose molds (Fig. SP1C right): prepare as many agarose molds as there are treatments using a negative rubber mold (Fig. SP1C left):

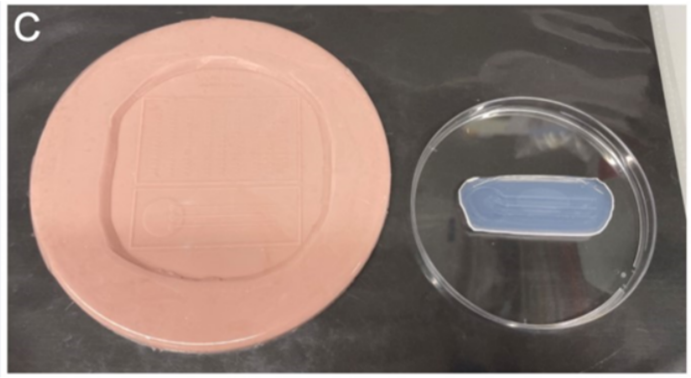

Agarose molds (Fig. SP1C right) a negative rubber mold (Fig. SP1C left)

4.1 Heat 1.5% agarose in 100 mL ddH2O in a beaker in a microwave in 00:00:30 bursts until the liquid is completely clear (no turbidity visible)

4.2 Carefully pour the heated agarose onto the rubber mold with channels from Ellett & Irimia [29] avoiding the formation of any bubbles.

4.3 Once it has solidified, peel the agarose mold off the rubber mold.

4.4 Store at 4 °C in a petri dish wrapped in parafilm until use.

Note

Heated agarose can be stored at 60 °C in between making several molds to prevent cooling/solidification.

#### Part 1: Zebrafish pre-experiment preparation (1 day post fertilization (dpf))

5 In the morning, move the zebrafish to a petri dish (LGG Labware, 90x16 mm, sterile) filled with about 20 mL E3 zebrafish water plus N-Phenylthiourea (PTU) (final PTU concentration: 0.003%) to avoid their pigmentation (see Table 1 on how to prepare E3 water and PTU).

6 In the evening, add 20 µL pronase (30 mg/mL) to the zebrafish (final pronase concentration: 0.03 mg/mL) and mix well to ensure that they will dechorionate (hatch from the chorion, which is the protective membrane around the zebrafish embryo) until the next morning.

#### Part 2: Preparation of bacterial suspensions (2 dpf)

7 Take fluorescently tagged bacterial species from 25% glycerol stocks that were stored at -80 °C and either streak them on an agar plate to pick a single colony (after overnight

growth at 37 °C ) for the following inoculation or directly inoculate them into 10 mL fresh LB medium.

8 Grow Overnight at 37 °C and 170 rpm with aeration until they reach the stationary phase.

- Depending on the biological question being asked, exponentially growing cells could also be used.

9 The next day, wash cells by centrifuging them at 7500 rcf, 00:03:00 . Discard the supernatant and replace it with 0.8% NaCl to resuspend the cell pellet.

10 Repeat step 9.

11 Measure the optical density at a wavelength of 600 nm and adjust it to reach the desired cell numbers per bacterial species (to be determined in a pre-experiment).

12 Mix 9 µL per bacterial culture or 0.8% NaCl solution (as control) with 1 µL of 0.5% phenol red (final concentration: 0.05%) to visualize whether an infection is successful and remains local in the inner ear structure, the otic vesicle, of the zebrafish.

> **Note**
>
> For multispecies infections, mix different bacterial species (e.g. in a 1:1 ratio) directly before loading the needle (i.e. before step 3 of part 3) to minimize interactions between them before reaching the host. Then proceed by adding 1 µL of 0.5% phenol red to the 9 µL mixed bacterial culture.

#### Part 3: Zebrafish infections into the otic vesicle (2 dpf)

13 Add approximately 9 mL of E3 zebrafish water with PTU (final PTU concentration: 0.003%) and 10 drops of the thawed anesthetic tricaine (ethyl 3-aminobenzoate methanesulfonate salt analytical standard, 4000 mg/L) using a Pasteur pipette (LLG Labware, 3 mL, unsterile) into a small petri dish (Greiner, 60x15mm, sterile).

14 Add 10-20 2 dpf zebrafish into this dish to anesthetize them (Fig. SP2A).

> **Note**
>
> Careful: try to add as little liquid as possible when transferring the zebrafish into the small petri dish to avoid diluting the tricaine concentration.

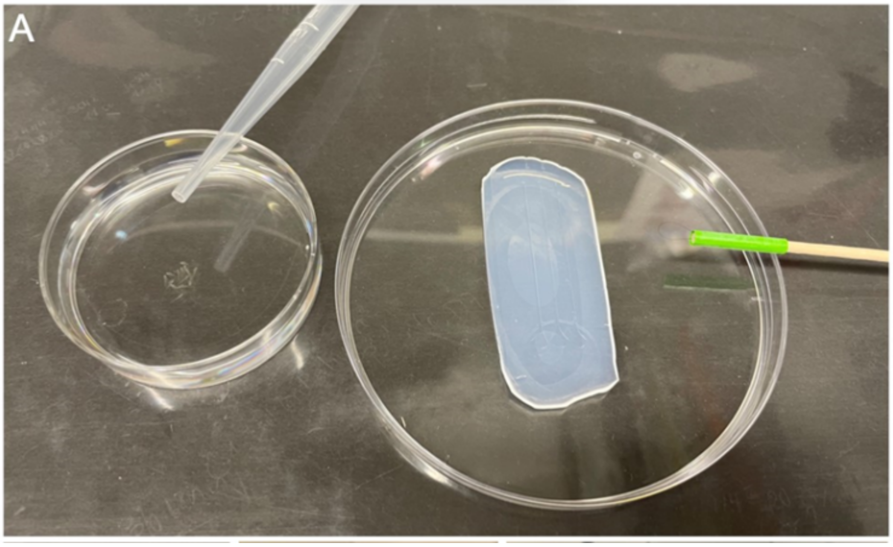

Figure SP2A. On the left are 2-days-postfertilization zebrafish in a small petri dish containing an anesthetizing solution (tricaine). A Pasteur pipette can be seen on top for moving the zebrafish from the petri dish onto the agarose mold. On the right is an agarose mold (placed on an inverted lid of a petri dish) and a hair loop manipulator.

15 For the first treatment, fill a glass needle (pulled borosilicate glass microcapillary injection needle (Science Products, 0.58mm diameter, GB100F-10)) with 10 µL of the 0.05% phenol red and 0.8% NaCl (control) or bacterial solution using a microloader tip (Eppendorf).

16 Put the filled glass needle into the micromanipulator, screwing it tight but not too tight to avoid breaking.

17 Position the needle in the field of view of the stereomicroscope using the micromanipulator.

18 Going from the lowest to the highest magnification, focus on the tip of the glass needle.

19 Once it is possible to distinguish the two walls of the needle at a final magnification of 56X, break off the tip of the glass needle using high-precision tweezers (e.g., RubisTech Switzerland, model 5-SA)(Fig. SP2C).

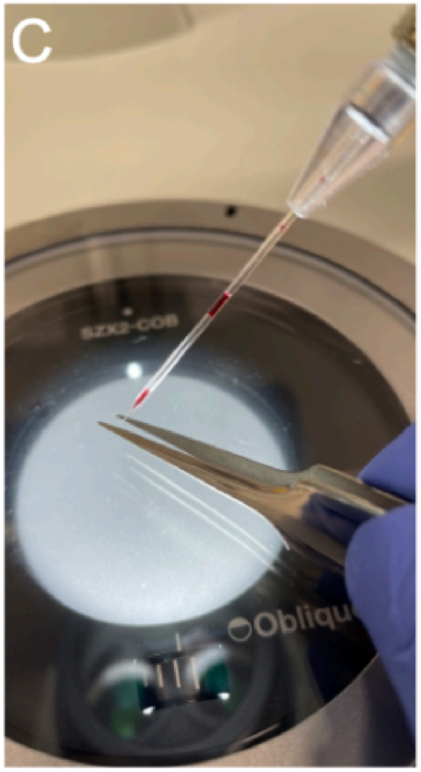

Figure SP2C. Breaking off the needle with fine tweezers using a stereomicroscope. The needle is held by a micromanipulator

20 Connect the injection pump with the micromanipulator (and thus the glass needle) via the tubing of the pump.

21 Put a drop of oil (VWR Chemicals, 10S) on a microscopy slide with a cavity (76x26 mm) that sits on an inverted lid of a petri dish (LGG Labware, 90x16 mm, sterile) and place it under the stereomicroscope.

22 Move the needle into the drop of oil using the micromanipulator and press 'Clean' on the injection pump to move the liquid into the tip of the needle.

23 Determine the desired drop size by injecting drops into the oil and adjusting either the injection time (to between 00:00:01 - 00:00:05 ) or injection pressure (to between 200-400 hPa). In case the solution in the needle gets aspirated back in or leaks, adjust the injection back pressure (to between 20-30 hPa). At a magnification of 56X, a drop that has the diameter of five stripes on the ruler corresponds to a volume of 1 nL. This can be determined because the liquid to inject will be a perfect sphere within the oil.

**Note**

Careful: the drop sinks to the bottom and will not be a perfect sphere anymore once it hits the bottom.

24 Whenever the needle is loaded and not in use, leave it in the oil to prevent evaporation of the liquid and the clogging of the needle.

25 Put a few drops of the E3-PTU-tricaine solution from the petri dish onto the agarose mold.

26 Make sure to wet the channels that will be used for the injection (lateral for the otic vesicle) using a hair loop manipulator.

27 Move the 2 dpf zebrafish from the small petri dish onto the large circle of the agarose mold for loading using a Pasteur pipette.

28 Load the zebrafish into the channels using a hair loop manipulator, turning all zebrafish onto the same side so that the same otic vesicle can be injected and imaged later on (left or right otic vesicle)(Fig. SP2B).

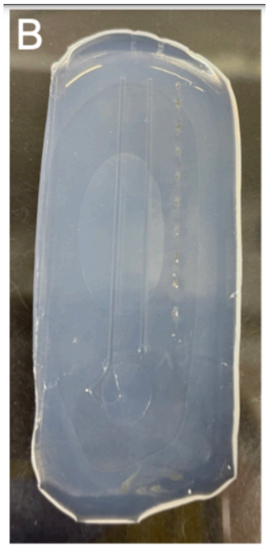

Figure SP2B. (B) An agarose mold with zebrafish aligned in the rightmost, the lateral channel.

29 Repeat step 14 to anesthetize the next batch of zebrafish for the same treatment. If it is a different treatment, this step is not necessary yet.

30 To determine the number of bacterial cells injected at the beginning of the injections, add a drop from the needle into 1 mL of 0.8% NaCl within a 1.5 mL Eppendorf tube.

31 Repeat step 30 to have at least duplicates for plating.

Note

Carefully move the glass needle into the NaCl solution of the Eppendorf tube without breaking it.

32 Place the mold under the objective of the stereomicroscope.

33 Using a magnification of 56X, position the needle into the otic vesicle and inject (Fig. SP2D).

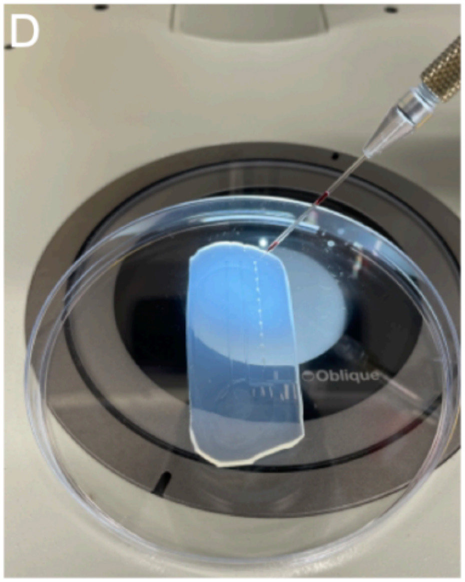

Figure SP2D. Injecting zebrafish using a stereomicroscope and a micromanipulator holding a glass needle.

Note

Careful: Make sure that the injection was applied correctly by checking whether the phenol red (bacterial) solution went into the otic vesicle and remained exclusively in there. Remove incorrectly injected zebrafish with the hair loop tool. Inject all zebrafish on the mold as described here.

34 Using a Pasteur pipette, aspirate the injected zebrafish and place them either individually into wells of a 48-well plate or five zebrafish individuals per well in a 24-well plate that is filled with E3 zebrafish water.

Note

Careful: first add some E3 zebrafish water onto the mold and then aspirate the zebrafish to prevent damaging them.

35 Repeat steps 27-34 if more zebrafish receiving the same treatment need to be injected.

36 To determine the number of bacterial cells injected at the end of the injections, add a drop from the needle into 1 mL of 0.8% NaCl within a 1.5 mL Eppendorf tube.

37 Repeat step 36 to have at least duplicates for plating.

**Note**

Carefully move the glass needle into the NaCl solution of the Eppendorf tube without breaking it.

38 Repeat steps 14-37 for each treatment.

39 Once all zebrafish from all treatments have been injected, check whether any zebrafish in the well-plate(s) have been damaged and remove them.

40 Incubate the well plate(s) at 28 °C either in the dark or with a 14:00:00 light and 10:00:00 dark cycle.

41 If necessary, further dilute and then plate 100 µL of the bacterial solution added to the 1 mL of 0.8% NaCl for all treatments in duplicates on 1.5% LB-agar plates to enumerate and confirm the infectious dose for each experiment.

**Note**

This step should also be used for pre-experiments to determine and define the number of cells to be injected per species.

42 Incubate plates at 37 °C (or the temperature required for your bacteria's growth) Overnight .

43 Count CFU from agar plates.

### Protocol (B): Zebrafish embedding and imaging (3 dpf)

In 1 collection

RESERVED DOI:
**10.17504/protocols.io.14egn6726l5d/v1**

Désirée A. Schmitz[1,2], Tobias Wechsler[1], Hongwei Bran Li[1,3], Bjoern H. Menze[1], Rolf Kümmerli[1]

[1]Department of Quantitative Biomedicine, University of Zurich, Zurich, Switzerland;
[2]Department of Microbiology, Harvard Medical School, Boston, Massachusetts, USA;
[3]Athinoula A. Martinos Center for Biomedical Imaging, Massachusetts General Hospital, Harvard Medical School, Boston, Massachusetts, USA

Desiree Schmitz
Harvard Medical School

**Protocol Info:** Désirée A. Schmitz, Tobias Wechsler, Hongwei Bran Li, Bjoern H. Menze, Rolf Kümmerli . Protocol (B): Zebrafish embedding and imaging (3 dpf). **protocols.io** **https://protocols.io/view/protocol-b-zebrafish-embedding-and-imaging-3-dpf-dc472yzn**

**Created:** April 16, 2024

**Last Modified:** May 01, 2024

**Protocol Integer ID:** 99199

#### Abstract

This protocol details the zebrafish embedding and imaging.

#### Guidelines

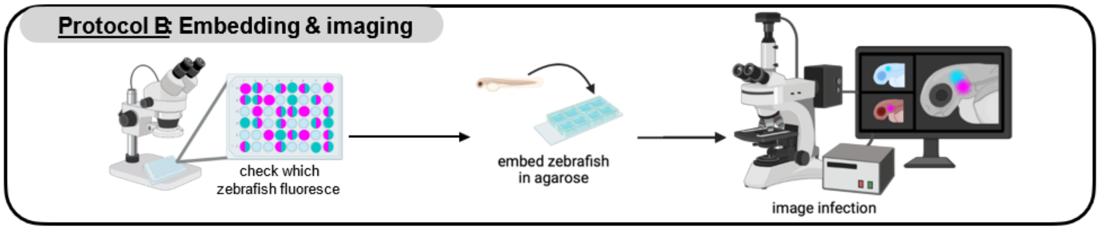

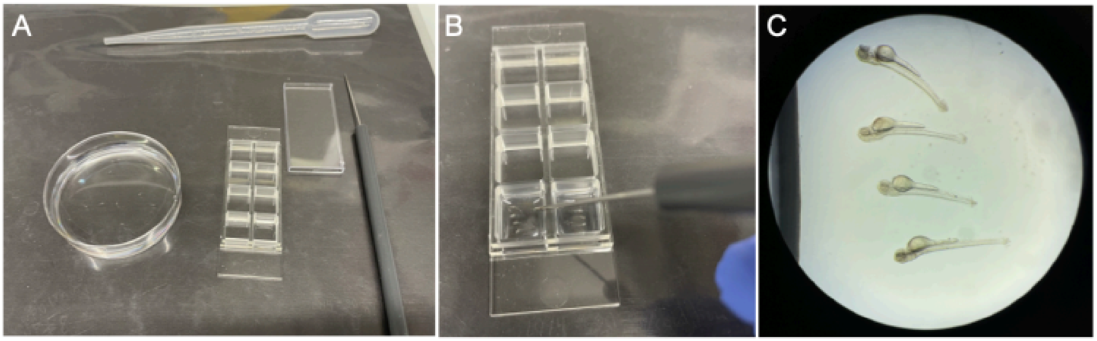

**Figure SP3. Embedding zebrafish.** (A) On the left are 3-days-post-fertilization (24 h post-infection) zebrafish in a small petri dish containing an anesthetizing solution (tricaine). Next to it is an Ibidi slide with the lid and a dissecting needle on its right side. On the top is a Pasteur pipette for moving zebrafish from the petri dish to the Ibidi µ slide. (B) Aligning zebrafish with a dissection needle in low-melting point agarose so that the infected otic vesicle is on the bottom. (C) View from a stereomicroscope to check whether zebrafish are properly aligned, facing the bottom with the otic vesicle thas has been infected (here: left otic vesicle).

#### Materials

**Equipment:**

For this protocol, you will need the following equipment:

- A stereomicroscope to embed zebrafish for imaging
- A heating block compatible with 1.5 mL Eppendorf tubes
- A widefield microscope with fluorescence filters that match the corresponding fluorophores of the tested bacterial species/ strains (e.g. Leica Thunder DMi8 widefield microscope with the Leica monochrome fluorescence DFC9000 GTC camera system)

#### Part 0: Material preparation

1 Prepare 1.5% low-melting point agarose by heating e.g. 1.5 g low-melting point agarose (Sigma, serial no: A9414) in 100 mL distilled water in a flask (minimal volume 150 mL) in 00:00:30 bursts in the microwave.

2 Once the solution is clear, i.e. no flocs or powder are visible anymore, aliquot into 1.5 mL Eppendorf tubes.

3 The day before embedding, prepare an Overnight culture of all bacteria used for injections. These will serve as a positive control.

#### Part 1: Embedding zebrafish in low-melting point agarose

4 Heat 1.5% low-melting point agarose in 1.5 mL Eppendorf tube aliquots to 90 °C in a heating block.

5 Once the agarose has melted, reduce the heat to 42 °C .

6 Check fluorescence in the otic vesicle of individual zebrafish using any microscope with sufficient fluorescence sensitivity and magnification. This step is only needed if the goal is to solely image zebrafish with an ongoing infection. These fishes can then be picked selectively based on their fluorescent signal in the otic vesicle.

7 Prepare a sufficiently large volume of anesthetic solution to use for all treatments: For this, add approximately 9 mL of E3 zebrafish water with PTU (final PTU concentration: 0.003%) and 10 drops of the thawed anesthetic tricaine (ethyl 3aminobenzoate methanesulfonate salt analytical standard, 4000 mg/L) using a Pasteur pipette (LLG Labware, 3 mL, unsterile) into a small petri dish (Greiner, 60x15mm, sterile).

8 For each treatment, move a few drops of the solution from the previous step into a separate small petri dish. This allows anesthetizing each treatment group individually to avoid cross-contamination.

9 Move the zebrafish of the first treatment into the anesthetic liquid in a designated petri dish (Fig. SP3A).

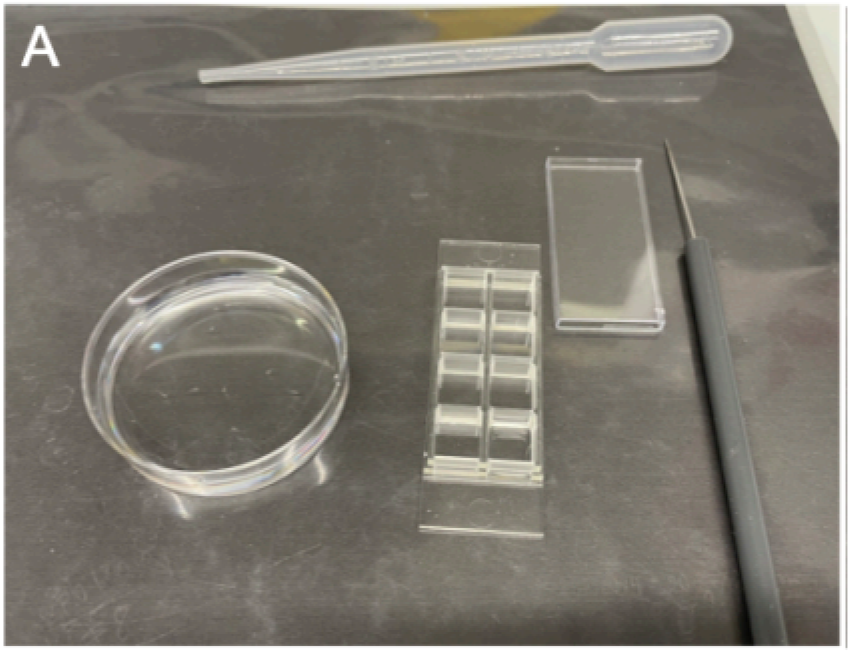

Figure SP3A. On the left are 3-days-post-fertilization (24 h postinfection) zebrafish in a small petri dish containing an anesthetizing solution (tricaine). Next to it is an Ibidi μ slide with the lid and a dissecting needle on its right side. On the top is a Pasteur pipette for moving zebrafish from the petri dish to the Ibidi μ slide.

Note

Careful: try to add as little E3 zebrafish water as possible with the zebrafish into the petri dishes to avoid diluting the concentration of the anesthetic.

10 Add 4-5 zebrafish of one treatment into an 8-well Ibidi μ slide (Vitaris, serial no: 80827-IBI) (Fig. SP3B).

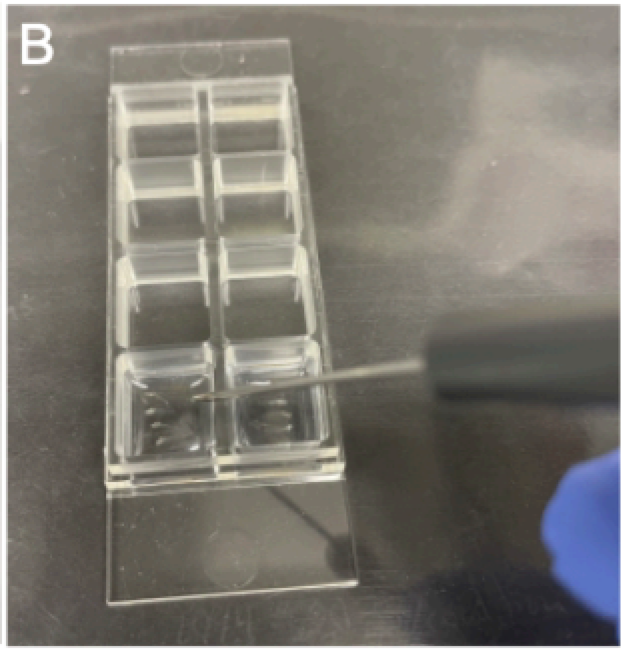

Figure SP3B. Aligning zebrafish with a dissection needle in low-melting point agarose so that the infected otic vesicle is on the bottom.

**Note**

Careful: again, try to add as little liquid as possible with the zebrafish into the well to avoid diluting the agar concentration.

11 Using a Pasteur pipette, add about 5 drops of the low-melting point agarose (from the 42 °C in the heating block) to the zebrafish in a well.

**Note**

The more agarose is added, the more time is usually required for positioning the zebrafish. However, the less agarose is added, the faster it will solidify.

12 Using a dissecting needle, mix the agarose and the small amount of liquid that was transferred with the zebrafish in the well so that the agarose concentration is the same throughout.

**Note**

Careful: do not damage the zebrafish with the needle.

13 Position the zebrafish with the dissecting needle so that the injected otic vesicle (see protocol A) is on the bottom of the slide to enable the use of an inverted widefield microscope for imaging (Fig. SP3C). Do this step with the naked eye as well as checking under a stereomicroscope.

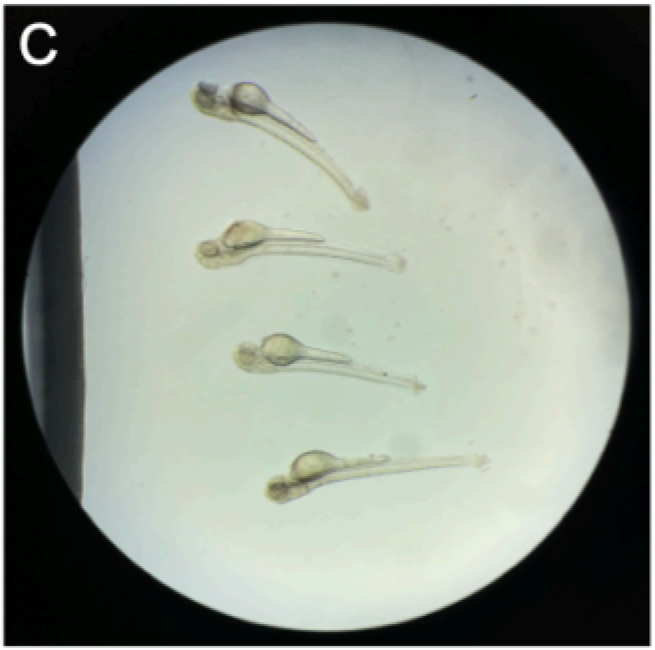

Figure SP3C. View from a stereomicroscope to check whether zebrafish are properly aligned, facing the bottom with the otic vesicle thas has been infected (here: left otic vesicle).

**Note**

Press the zebrafish onto the bottom of the well to reduce the working distance for the microscope, i.e., to improve the quality of the images.

14 Wait approximately 00:05:00 until the agarose has completely hardened.

15 Add E3 zebrafish water very carefully on top to ensure a constant supply of moisture.

16 Repeat steps 9-15 for all zebrafish that need to be embedded for imaging.

17 Add a positive control for each tagged bacterial species/strain, i.e., ~ 10 µL Overnight culture mixed with agarose into one well per species.

#### Part 2: Imaging the inner ear structure (otic vesicle) of zebrafish with a widefield microscope

18 Place up to 4 Ibidi slides into the slide holder in a widefield microscope.

19 Set up the brightfield and all required fluorescence channels, e.g., GFP (excitation at 475 nm & emission 520 nm), and mCherry (excitation at 555 nm & emission at 605 nm).

20 Set the positions of all otic vesicles with a 20X objective that has a long working distance (minimum 1 mm).

21 Image all saved positions of zebrafish otic vesicles from the previous step within the brightfield and fluorescence channels.

22 Save the images as .tiff files for further processing, also keeping the original images from the imaging software.

### Protocol (C): Automated segmentation of the otic vesicle and image analysis

In 1 collection

RESERVED DOI:
**10.17504/protocols.io.bp2l6219dgqe/v1**

Désirée A. Schmitz[1], Tobias Wechsler[1], Hongwei Bran Li[1,2], Bjoern Menze[1], Rolf Kümmerli[1]

[1]Department of Quantitative Biomedicine, University of Zurich, Zurich, Switzerland;
[2]Massachusetts General Hospital & Harvard Medical School, Harvard University, Boston, USA

Desiree Schmitz
Harvard Medical School

**Protocol Info:** Désirée A. Schmitz, Tobias Wechsler, Hongwei Bran Li, Bjoern Menze, Rolf Kümmerli . Protocol (C): Automated segmentation of the otic vesicle and image analysis. **protocols.io** **https://protocols.io/view/protocol-c-automated-segmentation-of-the-otic-vesi-dc482yzw**

**Created:** April 16, 2024

**Last Modified:** May 01, 2024

**Protocol Integer ID:** 99200

#### Abstract

This protocol details the automated segmentation of the otic vesicle and image analysis.

#### Materials

For this protocol, you will need the following scripts and software:

- Docker and the docker image (docker pull branhongweili/dqbm_cell_seg:v3.1) to run the pre-trained segmentation model.
- Depending on the operating system, either use the segmentation.sh shell script (for MacOS/Ubuntu) or the segmentation.ps1 PowerShell script (for Windows).
- An installation of FIJI.
- The check_measure.py script.
- An installation of R.
- The co_localization.R script.

#### Automated segmentation of the otic vesicle and image analysis

1

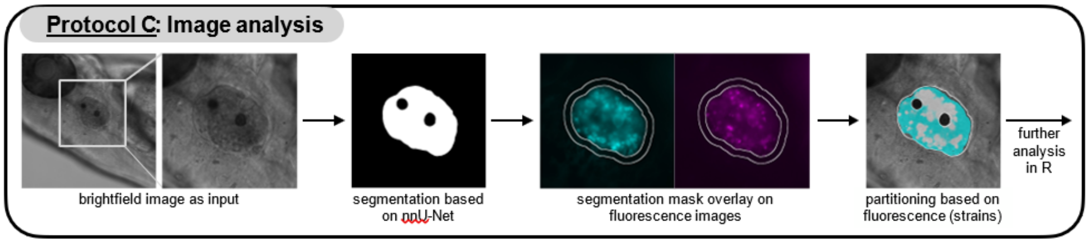

2 Organize your images in a folder with two subfolders:

- one for the brightfield images named ‘Images’,
- one for all fluorescence images named ‘Fluor’.

The image file names have to contain the slice number with a Z as a prefix and the channel number (0-based) with a C as a prefix (e.g., PositionName_Z00_C00.tif). For example, for measuring two different fluorophores and the brightfield image, we have three channels, C00 (e.g., brightfield), C01 (e.g., GFP), C02 (e.g., mCherry). We include an example script to convert Leica image files accordingly (sort_images_lif.py), which can be used as an example of how to organize all images automatically as described.

3 For the segmentation of the otic vesicle, run the segmentation.sh (segmentation.ps1 for Windows) script with the previously created directory as input (e.g. for Mac: in Terminal). Make sure the shell script is executable (e.g. for Mac: by typing

```
chmod +x segmentation.sh
```

in Terminal). The script will start a docker container and the segmentation model on the specified brightfield images.

4 In the same directory that contains your brightfield (Images) and fluorescence (Fluor) images, a directory named output_real_value is created that contains the segmentation masks.

5 Start FIJI and open the check_measure.py script.

6 Run the script and select the directory with your brightfield (Images) and fluorescence (Fluor) images and segmentation masks (output_real_value) as input.

7 FIJI will display the brightfield images and the corresponding segmentation. In the ROI Manager window untick the box “Show All” and select one of the ROIs in the list to only show the segmentation for a single ROI. Go through the ROI sequence and delete inaccurate segmentations.

8 Complete the manual review of the segmentations by pressing OK.

9 The script creates six files:

9.1 an image with an overlay of the strain partitioning based on fluorescence (e.g. Sample1_overlay.tif),

9.2 a text file containing the log to check what has been done up until here (e.g. Sample1_log.txt)

9.3 a zip file with the ROIs (Sample1_rois.zip)

9.4 a table with fluorescence values within the whole otic vesicle (mean and integrated density for both GFP and mCherry; e.g. Sample1 ov_data.csv)

9.5 A table with the size of the area occupied by tagged strains (e.g. Sample1_strain_count.csv) → e.g. needed to calculate co-localization

9.6 A table with the fluorescence values of individual pixels within the otic vesicle (e.g. Sample1 pixel_data.csv).

**Note**

Since the zebrafish shows inherent auto-fluorescence, we introduced a threshold value to delineate bacterial occupation from auto-fluorescence. The threshold value for bacterial occupation is defined as twice the fluorescence value observed in the surrounding tissue of the otic vesicle (a layer of approximately 16 µm, corresponding to 50 pixels).

10 Use the accompanying R-script to visualize the size of the area occupied by each tagged bacterial species and the overlap between the two species to quantify their co-localization (script: co_localization.R).

11 Check whether the fluorescence images in FIJI correspond to the values in R by dropping the Fluor folder into FIJI and comparing it to the R plot.
