## Supplementary material for "A new protocol for multispecies bacterial infections in zebrafish and their monitoring through automated image analysis": S2 File Supplementary Figures

**Supplementary Information**
This file contains supplementary information for the article

### A new protocol for multispecies bacterial infections in zebrafish and their monitoring through automated image analysis

Désirée A. Schmitz[12], Tobias Wechsler[1], Hongwei Bran Li[13], Bjoern H. Menze [1] & Rolf Kümmerli [1]

[1] Department of Quantitative Biomedicine, University of Zurich, Zurich, Switzerland
[2] Department of Microbiology, Harvard Medical School, Boston, Massachusetts, USA
[3] Athinoula A. Martinos Center for Biomedical Imaging, Massachusetts General Hospital, Harvard Medical School, Boston, Massachusetts, USA

The supplementary information comprises:

- 3 supplementary Figures: Figure S1 – S3

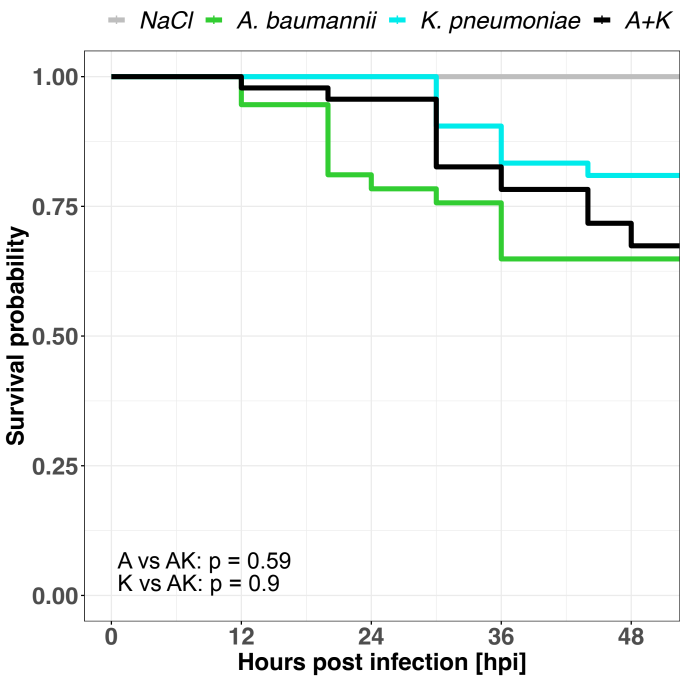


**Figure S1. Kaplan Meier survival curves of 2 days post-fertilization zebrafish larvae with mono- and co-infections.** Survival of zebrafish with co-infections (black curve) and the corresponding mono-infections (colored curves). The mean number of CFU is 6850 for *K. pneumoniae* and 8600 for *A. baumannii.* Survival (y-axis) was monitored every 4-8 h for 54 h in total (x-axis). Control larvae were injected with a 0.8% NaCl solution. We conducted multiple pairwise comparisons using the log-rank test (alpha = 0.05, Benjamini-Hochberg p-value adjustment). Data are from two independent experiments with a total of 37-53 individual zebrafish per treatment.

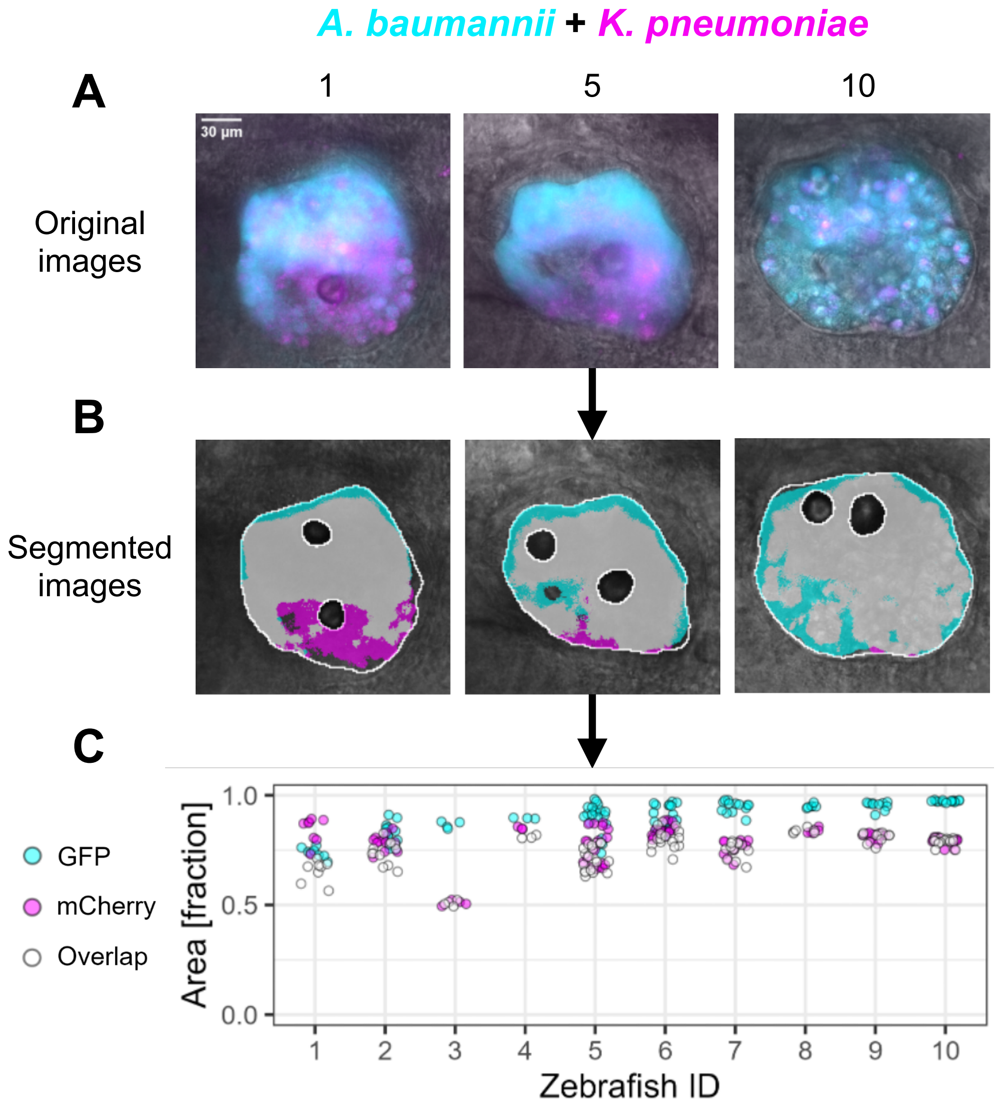


**Figure S2. The two co-infecting species *A. baumannii* and *K. pneumoniae* can reliably be detected and quantified in the otic vesicle of zebrafish.** (A) Representative images of three individual zebrafish are shown as overlays of the brightfield, GFP (excitation at 475 nm & emission at 520 nm; shown in cyan), and mCherry (excitation at 555 nm & emission at 605 nm; shown in magenta) image. (B) Masks obtained from automated segmentation are shown in white. The partitioning of the bacterial strains is done using the fluorescence signal. Since the zebrafish show some inherent auto-fluorescence, bacterial occupation is defined by a fluorescence value that is twice as high as the fluorescence value observed in the surrounding tissue of the otic vesicle (a layer of approximately 16 µm, corresponding to 50 pixels). (C) Quantitative image analysis showing the relative area of the otic vesicle that is occupied by the co-infecting pathogens (y-axis) across zebrafish individuals (x-axis), ordered from lowest to highest bacterial occupation. Magenta and cyan fractions respectively represent the area occupied by either *K. pneumoniae* or *A. baumannii*, with the mean number of CFU injected being 8100 and 7400, respectively. The grey fraction (open circles) represents the area simultaneously occupied by both pathogens. Each data point represents a z-slice imaged from the respective zebrafish ID. The data shown are from one experiment with n=10.

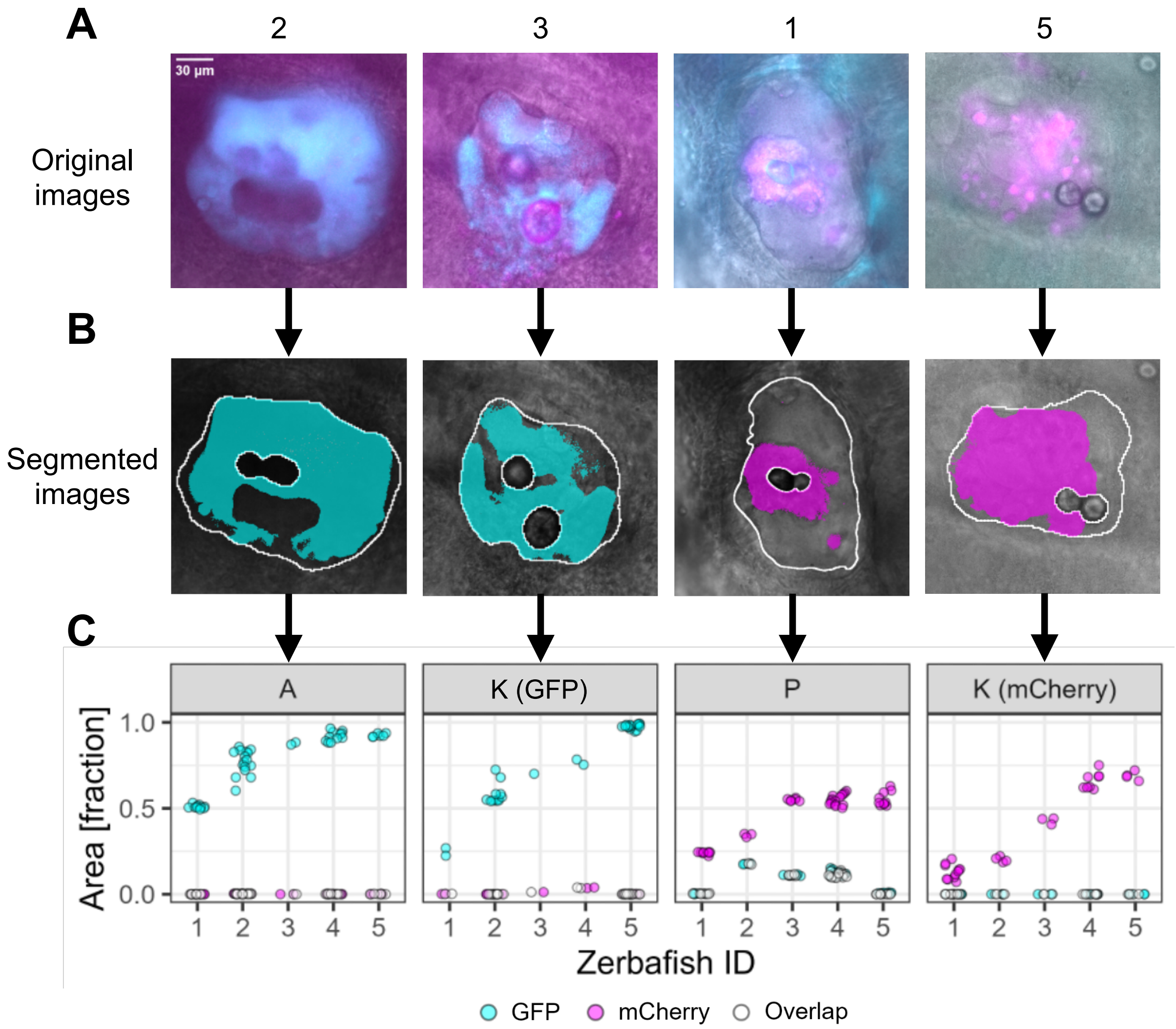


**Figure S3. A mono-infection can reliably be detected and quantified in the otic vesicle of zebrafish.** (A) Representative images of individual zebrafish per pathogen are shown as overlays of the brightfield, GFP (excitation at 475 nm & emission at 520 nm; shown in cyan), and mCherry (excitation at 555 nm & emission at 605 nm; shown in magenta) image. (B) Masks obtained from automated segmentation are shown in white. Since the zebrafish show some inherent auto-fluorescence, bacterial occupation is defined by a fluorescence value that is twice as high as the fluorescence value observed in the surrounding tissue of the otic vesicle (a layer of approximately 16 µm, corresponding to 50 pixels). (C) Quantitative image analysis showing the relative area of the otic vesicle that is occupied by a pathogen (y-axis) across zebrafish individuals (x-axis), ordered from lowest to highest bacterial occupation. The GFP signal is shown in cyan (*A. baumannii* and *K. pneumoniae* on left) and the mCherry signal in magenta (*P. aeruginosa* and *K. pneumoniae* on right). The mean number of CFU injected from left to right is as follows: 7400, 9200, 13200, and 8100. The grey fraction (open circles) represents the area when both signals overlap. Each data point represents a z-slice imaged from the respective zebrafish ID. The data shown are from two individual experiments, with n=5 for each mono-infection.
