## Supplementary material for "A new protocol for multispecies bacterial infections in zebrafish and their monitoring through automated image analysis": S4 File Image analysis scripts: check_measure.py

```
# title				: check_measure.py
# description		: create ROIs from segmentation, select ROIs and measure expression, pixels, strain area 
# author			: Tobias Wechsler
# date				: 2023/09/14
# Notes				: for use with ImageJ and Jython (Python in Java)
# ======================================================================================================================
#
# USAGE:
# This script requires images to be organized in a folder with three subfolders: 
# "Images" for bright field images, "Fluor" for fluorescence images, and "output_real_value" for segmentation images.
# The image file names must contain "Z[slice number]" and "C[channel number with one leading 0]",
# indicating the slice and channel respectively. Segmentation images should be in PNG format, others in TIFF.
#
# INPUT FOLDER STRUCTURE:
#
# Sample001
#	└── Images 
#	│		├── Sample001_C00_Z00.tif
#	│		├── Sample001_C00_Z01.tif
#	│		└── ...
#	│
#	├── Fluor
#	│		├── Sample001_C01_Z00.tif
#	│		├── Sample001_C02_Z00.tif
#	│		└── ...
#	│
#	└── output_real_value
#			├── Sample001_C00_Z00.png
#			├── Sample001_C00_Z01.png
#			└── ...
	

# Import required libraries
import os
import re
import csv
import sys
from datetime import datetime
from ij import IJ, ImageStack, ImagePlus
from ij.plugin import FolderOpener, Duplicator, ImageCalculator, RoiEnlarger
from ij.plugin.frame import RoiManager
from ij.gui import ShapeRoi, WaitForUserDialog
from ij.measure import ResultsTable

# Input: Selection of input folder through ImageJ interface
#@ File (label="Select an input folder", style="directory") IN_DIR

# Functions:

def get_info(filename):
	'''
	Given a filename, returns the channel and slice number
	'''

	ch = int(re.search('C\d+', filename).group().replace('C',''))
	z = int(re.search('Z\d+', filename).group().replace('Z',''))

	return ch, z

def log_input_folder(in_dir):
	'''
	Writes a log with the contents of the selected input folder.
	'''

	IJ.log(os.path.basename(in_dir))
	for d, sd, fs in os.walk(in_dir):
		if d!= in_dir:
			IJ.log('   |--{0}'.format(os.path.basename(d)))
			for f in sorted(fs):
				IJ.log('   |     |-- {0}'.format(f))
			IJ.log('   |')
	IJ.log('')

def make_roi(seg):
	'''
	Create a ROI for the OV from the segmenataion, where OV has the value 1 and CD (black dots)  2,
	and the background ROI excluding the OV
	'''

	seg_name = seg.getTitle()
	back_roi = get_background(seg)


	IJ.setRawThreshold(seg, 1, 1)
	IJ.run(seg, "Analyze Particles...", "clear include add")
	RM = RoiManager.getInstance()
	rois = RM.getRoisAsArray()
	
	
	if len(rois) > 0:
		# select biggest ROI if there are more then
		max_size = 0
		max_roi_index = 0
		for j, roi in enumerate(rois):
			if roi.size() > max_size:
				max_size = roi.size()
				max_roi_index = j

		ov_roi = ShapeRoi(rois[max_roi_index])
		
		# remove CD (black dots) from ov_roi
		IJ.setRawThreshold(seg, 2, 2)
		IJ.run(seg, "Analyze Particles...", "clear include add")
		
		cd_rois = RM.getRoisAsArray()
		for cd in cd_rois:
			ov_roi = ov_roi.not(ShapeRoi(cd))
		
		RM.reset()
		ov_roi.setName(seg_name)

		return ov_roi, back_roi 


def get_background(seg):
	'''
	Create a ROI around the OV to get a local background fluorescence value
	'''

	IJ.setRawThreshold(seg, 1, 2)
	IJ.run(seg, "Analyze Particles...", "clear include add")
	RM = RoiManager.getInstance()
	rois = RM.getRoisAsArray()
	
	# get biggest roi, maybe better to combine them
	if len(rois) > 0:
		max_size = 0
		max_roi_index = 0
		for j, roi in enumerate(rois):
			if roi.size() > max_size:
				max_size = roi.size()
				max_roi_index = j
		
		ov_roi = rois[max_roi_index]
		big_roi = ShapeRoi(RoiEnlarger().enlarge(ov_roi, 50))
		near_roi = big_roi.xor(ShapeRoi(ov_roi))

		return near_roi

def select_ROI(rois, img):
    '''
    Allows manual selection and removal of bad ROIs using the ROI Manager.
    '''

    RM = RoiManager.getInstance()
    if not RM:
        RM = RoiManager()
    RM.reset()

    for roi in rois:
    	RM.addRoi(roi)


    RM.runCommand(img, "Show All without labels")
    WaitForUserDialog('Delete bad ROIs in the RoiManager').show()

    new_rois = RM.getRoisAsArray()
    z_positions = [r.getPosition() for r in new_rois]
    RM.reset()

    return new_rois, z_positions


def seg_strains(fluor, roi, back_roi):
	'''
	Create images showing the area occupied by a strain within the OV.
	The threshold is set to double the mean expression in the back ground

	'''

	IJ.run(fluor, "Select None", "")
	RM = RoiManager.getInstance()

	back_roi.setImage(fluor)
	average = float(back_roi.getStatistics().mean)
	# set threshold (default: double of the mean background)
	low_threshold = 2 * average
	RM.reset()
	RM.addRoi(roi.getInverse(fluor))
	RM.select(fluor, 0)
	IJ.run(fluor, "Clear", "slice")
	IJ.setRawThreshold(fluor, low_threshold , 65535)
	RM.reset()
	
	IJ.run(fluor, "Convert to Mask", "")

	return fluor, average


def measure_pixels(img, roi):
	'''
	Retrieves fluorescence values from individual pixels within a ROI.
	'''
	x = []
	y = []
	val = []
	points = roi.getContainedPoints()
	for p in points:
		x.append(p.x)
		y.append(p.y)
		val.append(img.getPixel(p.x, p.y)[0])

	return x, y, val


if __name__ == '__main__':
	IJ.run("Close All", "")
	IJ.log(datetime.now().isoformat())

	# Setup input folders
	pos_name = IN_DIR.getName()
	pos_path = IN_DIR.getPath()
	in_dir = os.path.join(pos_path, 'Images')
	fluor_dir = os.path.join(pos_path, 'Fluor')
	seg_dir = os.path.join(pos_path, 'output_real_value')

	log_input_folder(pos_path)
	
	# Make file lists
	bf_imgs = sorted([f for f in os.listdir(os.path.join(in_dir)) if f.endswith('.tif')])
	fluor_imgs = sorted([f for f in os.listdir(os.path.join(fluor_dir)) if f.endswith('.tif')])
	seg_imgs = sorted([f for f in os.listdir(os.path.join(seg_dir)) if f.endswith('.png')])

	# Check for present channels
	channels = []
	zslices = []
	for img in fluor_imgs:
		ch ,z = get_info(img)
		channels.append(ch)
		zslices.append(z)
	zslices = [z - min(zslices) for z in zslices]
	channels = set(channels)
	zslices = set(zslices)
	IJ.log('Fluorescence channels: {0}'.format(channels))
	IJ.log('Slices (0 based): {0}'.format(zslices))


	# Check which slices have segmentation
	IJ.log('\nImages    :       Segmentation\n=============================================')
	offset = 0
	z_positions = []
	for i, img in enumerate(bf_imgs):
		ch, z = get_info(img)
		if len(seg_imgs) >= i-offset+1:
			seg_img = seg_imgs[i - offset]
			l_ch, l_z = get_info(seg_img)
			if z == l_z:
				IJ.log('{0} : {1}'.format(img, seg_img))
				z_positions.append(i + 1)
			else:
				IJ.log('{0} : -'.format(img))
				offset += 1
		else:
			IJ.log('{0} : -'.format(img))

	IJ.log('Segmented Slices (1 based): {0}'.format(z_positions))

	# Make ROIs from segmentation
	
	IJ.log('\nCreate ROIs...\n')
	ov_rois = []
	back_rois = []

	for i, img_file in enumerate(seg_imgs):
		seg = IJ.openImage(os.path.join(seg_dir, img_file))
		ov_roi, back_roi = make_roi(seg)
		if ov_roi.getLength() > 0:
			ov_roi.setPosition(z_positions[i])
			back_roi.setPosition(z_positions[i])
			ov_rois.append(ov_roi)
			back_rois.append(back_roi)
			IJ.log('{0}   :   OK'.format(img_file))
		else:
			IJ.log('{0}   :   FAILED'.format(img_file))
		seg.close()
	
	RM = RoiManager.getInstance()
	if not RM:
		RM = RoiManager(True)
	RM.reset()

	# Manually check the ROIs

	IJ.log('\nCheck ROIs...\n')
	bfield = FolderOpener.open(in_dir)
	bfield.setTitle('Brightgield')
	bfield.show()
	IJ.run(bfield, "Grays", "")


	temp_rois, z_positions = select_ROI(ov_rois, bfield)
	back_rois = [r for r in back_rois if r.getPosition() in z_positions]
	for roi in ov_rois:
		if roi.getPosition() in z_positions:
			IJ.log('{0}   :   OK'.format(roi.getName()))
		else:
			IJ.log('{0}   :   EXCLUDED'.format(roi.getName()))
			
	
	ov_rois = temp_rois

	n_good_rois = len(ov_rois)

	if n_good_rois == 0:
		quit()
	for roi in ov_rois:
		RM.addRoi(roi)

	bfield.hide()

	# Measure ROIs
	
	IJ.log('\nMeasure ROIs...\n')
	fluorC1 = FolderOpener.open(fluor_dir, " filter=C01")
	fluorC1.setTitle("GFP")
	IJ.run(fluorC1, "Select None", "")
	RM.runCommand(fluorC1, "Measure")

	IJ.log('Measured {0} ROIs in C01'.format(RM.getCount()))
	if 2 in channels:
		fluorC2 = FolderOpener.open(fluor_dir, " filter=C02")
		IJ.run(fluorC2, "Select None", "")
		fluorC2.setTitle("mCherry")
		RM.runCommand(fluorC2, "Measure")
		IJ.log('Measured {0} ROIs in C02'.format(RM.getCount()))

	ResTab = ResultsTable.getResultsTable()

	for i in range(ResTab.size()):
		ResTab.setValue('Sample', i , pos_name)

	ResTab.save(os.path.join(pos_path, '{0}_ov_data.csv'.format(pos_name)))
	ResTab.reset()

	# Segment strains

	IJ.log('\nMeasure strain segments...\n')
	RM.reset()

	gfp_segs = []
	mch_segs = []
	gfp_back = []
	mch_back = []

	offset = 0

	for i in zslices:
		z = i + 1
		if z in z_positions:

			IJ.run(fluorC1, "Select None", "")
			img_c1 = Duplicator().run(fluorC1, 1,1,z,z,1,1)
			gfp_seg, gfp_back_val  = seg_strains(img_c1, ov_rois[i-offset], back_rois[i-offset])
			img_c1.close()

			gfp_segs.append(gfp_seg)
			gfp_back.append(gfp_back_val)

			if 2 in channels:
				IJ.run(fluorC2, "Select None", "")
				img_c2 = Duplicator().run(fluorC2, 1,1,z,z,1,1)
				mch_seg, mch_back_val = seg_strains(img_c2, ov_rois[i-offset], back_rois[i-offset])
				img_c2.close()

				mch_segs.append(mch_seg)
				mch_back.append(mch_back_val)
				IJ.log('Slice {0}   :  Segmented C01 and C02'.format(i + 1))
			else:
				IJ.log('Slice {0}   :  Segmented C01'.format(i + 1))
		else:
			IJ.log('Slice {0}   :  Not Segmented'.format(i + 1))
			offset +=1
			IJ.run(fluorC1, "Select None", "")
			img_c1 = Duplicator().run(fluorC1, 1,1,1,1,1,1)
			IJ.run(img_c1, "Set...", "value=0")
			gfp_segs.append(img_c1)
			if 2 in channels:
				IJ.run(fluorC2, "Select None", "")
				img_c2 = Duplicator().run(fluorC2, 1,1,1,1,1,1)
				IJ.run(img_c2, "Set...", "value=0")
				mch_segs.append(img_c2)

	ResTab = ResultsTable.getResultsTable()
	ResTab.reset()
	RM.reset()

	# Measure strain segments

	IJ.run("Set Measurements...", "area mean integrated stack display redirect=None decimal=3")

	for roi in ov_rois:
		RM.addRoi(roi)

	gfp_stack = ImagePlus('GFP_tag', ImageStack.create(gfp_segs))

	gfp_stack.show()
	IJ.run(gfp_stack, "Divide...", "value=255 stack")
	RM.runCommand(gfp_stack,"Deselect")
	RM.runCommand(gfp_stack,"Measure")

	if 2 in channels:
		mch_stack = ImagePlus('MCH_tag', ImageStack.create(mch_segs))
		

		mch_stack.show()
		IJ.run(mch_stack, "Divide...", "value=255 stack")
		RM.runCommand(mch_stack,"Deselect")
		RM.runCommand(mch_stack,"Measure")
		#IJ.run(ImagePlus(), "Merge Channels...", "c1=MCH_tag c2=GFP_tag create")

		overlap_stack = ImageCalculator().run(gfp_stack, mch_stack, 'and stack create')
		RM.runCommand(overlap_stack,"Deselect")
		RM.runCommand(overlap_stack,"Measure")
		overlap_stack.close()

		IJ.run(ImagePlus(), "Merge Channels...", "c1=MCH_tag c2=GFP_tag create")
	else:
		IJ.run(gfp_stack, "Green","")


	ResTab = ResultsTable.getResultsTable()

	ResTab.renameColumn("Mean", "Strain_Area_fraction")
	ResTab.renameColumn("RawIntDen", "Strain_Area")

	for i in range(ResTab.size()):
		ResTab.setValue('Sample', i , pos_name)

	ResTab.save(os.path.join(pos_path, '{0}_strain_count.csv'.format(pos_name)))

	strain_img = IJ.getImage()

	IJ.run(strain_img, "Multiply...", "value=255.0")
	
	strain_img.setC(2)
	IJ.run(strain_img, "Cyan", "")
	strain_img.setC(1)
	IJ.run(strain_img, "Magenta", "")
		


	strain_img.hide()

	# Measure pixels

	IJ.log('\nMeasure pixels...\n')
	pixel_data = {'Sample':[],
			'x':[],
			'y':[],
			'Slice':[],
			'GFP_C01':[],
			'MCH_C02':[]}

	for roi in ov_rois:
		z = roi.getZPosition()
		IJ.run(fluorC1, "Select None", "")
		img_c1 = Duplicator().run(fluorC1, 1,1,z,z,1,1)
		x, y, gfp = measure_pixels(img_c1, roi)
		img_c1.close()
		if 2 in channels:
			IJ.run(fluorC2, "Select None", "")
			img_c2 = Duplicator().run(fluorC2, 1,1,z,z,1,1)
			_, _, mch = measure_pixels(img_c2, roi)
			img_c2.close()
			
		pixel_data['x'].extend(x)
		pixel_data['y'].extend(y)
		pixel_data['GFP_C01'].extend(gfp)
		if 2 in channels:
			pixel_data['MCH_C02'].extend(mch)
		else:
			pixel_data['MCH_C02'].extend(['NA']*len(x))
		pixel_data['Slice'].extend([z]*len(x))
		pixel_data['Sample'].extend([pos_name]*len(x))
		IJ.log('{0}   :  {1} pixels'.format(roi.getName(), len(gfp)))
		

	with open(os.path.join(pos_path,'{0}_pixel_data.csv'.format(pos_name)), 'wb') as f:
		writer = csv.writer(f)
		writer.writerow(['Sample','x','y','Slice','GFP_C01','MCH_C02'])
		for i in range(len(pixel_data['Sample'])):
			writer.writerow([pixel_data['Sample'][i],
								pixel_data['x'][i],
								pixel_data['y'][i],
								pixel_data['Slice'][i],
								pixel_data['GFP_C01'][i],
								pixel_data['MCH_C02'][i]])

 	# Make overlay image

	bfield.show()
	IJ.run(bfield, "Grays", "")

	over_img = bfield
	over_img.setTitle('Overlay')
	RM.runCommand("Save", os.path.join(pos_path, pos_name + '_rois.zip'))
	RM.runCommand(over_img, "Show All without labels")
	IJ.run("From ROI Manager", "")

	strain_img.setTitle('strains')
	if 2 in channels:
		IJ.run(strain_img, "Stack to RGB", "slices")
	else:
		IJ.run(strain_img, "RGB Color", "")

	for i in range(over_img.getNSlices()):
		olay = Duplicator().run(strain_img, 1,1,i+1,i+1,1,1)
		olay.setTitle('strainol')
		olay.show()
		over_img.setSlice(i+1)
		
		IJ.selectWindow(over_img.getTitle())

		IJ.run("Add Image...", "image=strainol x=0 y=0 opacity=60 zero")
		olay.close()


	out_path = os.path.join(pos_path, pos_name + '_overlay.tif')
	IJ.saveAs(over_img, "Tiff", out_path)
	

	IJ.log('Finished {0}'.format(datetime.now().isoformat()))
	IJ.selectWindow("Log")
	IJ.saveAs("Text", os.path.join(pos_path, pos_name + "_log.txt"))

	IJ.log('### LOG SAVED: {0} ###'.format(os.path.join(pos_path, pos_name + "_log.txt")))
	print('Finished {0}'.format(datetime.now().isoformat()))
```
