## Supplementary material for "A new protocol for multispecies bacterial infections in zebrafish and their monitoring through automated image analysis": S4 File Image analysis scripts: co_localization.R

```
# This script is designed for analyzing and visualizing the area within OVs occupied by bacteria

# Load necessary libraries
library(ggplot2)
library(dplyr)
library(tidyr)
library(ggpubr)

# List CSV files 

# Set the directory path containing all sample directories
in_dir = '20231116_AKP'

# List all files in the directory that match the '_strain_count.csv' pattern
stn_files <- list.files(path=in_dir, pattern = '_strain_count.csv$', recursive = T, full.names = T)

# Read and combine data from all listed CSV files
stn_tables <- lapply(stn_files, read.csv)
stn_data <- bind_rows(stn_tables)
rm(stn_tables)


# Create GFP, MCH and overlap tag
stn_data$tag <-substr(stn_data$Label, 1,3) 
stn_data[which(stn_data$tag=='Res'), 'tag'] <- 'Overlap'
stn_data[which(stn_data$tag=='MCH'), 'tag'] <- 'mCherry'

# Create a jitter plot to visualize the area fraction of strains across samples
area.frc <- ggplot(stn_data, aes(x=Sample, y=Strain_Area_fraction, fill = tag)) +
              geom_jitter(shape = 21, width = 0.2, alpha = 0.5, size = 2) +
              scale_fill_manual(values = c('cyan', 'magenta', '#FFFFFF')) +
              theme_bw(base_size = 18) + 
              theme(legend.position = 'bottom', legend.title = element_blank(), legend.margin = margin(t=0.0, unit = 'cm')) +
              scale_x_discrete(labels = seq(1,10)) +
              scale_y_continuous(breaks = c(0, 0.5, 1), limits = c(-0.001,1)) +
              ylab('Area [fraction]')+ xlab('Zebrafish ID');area.frc
```
