## Supplementary material for "A new protocol for multispecies bacterial infections in zebrafish and their monitoring through automated image analysis": S4 File Image analysis scripts: segmentation.ps1

```
# Check if a path is provided
param(
    [string]$path
)

if (-not $path) {
    Write-Host "No path provided. Please provide a folder path."
    exit 1
}

# Image folder path
$INPUT_FOLDER = Join-Path -Path $path -ChildPath "Images"


# Run Docker container
$CONTAINERID = docker run -dit -e NVIDIA_DISABLE_REQUIRE=true `
    -v "${INPUT_FOLDER}:/app/input:ro" `
    --env 'nnUNet_raw=/app/nnUNet_raw' `
    --env 'nnUNet_preprocessed=/app/nnUNet_preprocessed' `
    --env 'nnUNet_results=/app/nnUNet_results' branhongweili/dqbm_cell_seg:v3.1

# Pre-processing
docker exec -t $CONTAINERID python3 pre.py

# Segmentation with u-net
docker exec -t $CONTAINERID nnUNetv2_predict -i /app/input_pre/$SAMPLE_NAME -o /output -d 002 -c 2d -f all -device cpu

# Post-processing It upsamples the masks from 512*512 to 2048*2048 and save an overlay version to check the segmentation quality. 
# If the original image size is not 2048*2048, you could add ``--size {YOUR_SIZE}` to resize it to the original size you wish.   
docker exec -t $CONTAINERID python3 post.py --alpha 0.20 

# Copy the segmentation to the output folder
docker cp "${CONTAINERID}:/output_real_value/" $path

# Stop and remove the Docker container
docker stop $CONTAINERID
docker rm $CONTAINERID
```
