## Supplementary material for "A new protocol for multispecies bacterial infections in zebrafish and their monitoring through automated image analysis": S4 File Image analysis scripts: segmentation.sh

```
#!/bin/bash

# Check if a path is provided
if [ $# -eq 0 ]; then
    echo "No path provided. Please provide a folder path."
    exit 1
fi


# Convert to absolute path
ABS_PATH=$(realpath "$1")

# Image folder path
INPUT_FOLDER="$ABS_PATH/Images"

# Run Docker container
CONTAINERID=` docker run -dit -e NVIDIA_DISABLE_REQUIRE=true \
         -v $INPUT_FOLDER:/app/input:ro  \
         --env 'nnUNet_raw=/app/nnUNet_raw' \
         --env 'nnUNet_preprocessed=/app/nnUNet_preprocessed' \
         --env 'nnUNet_results=/app/nnUNet_results' branhongweili/dqbm_cell_seg:v3.1 `

# Sre-processing
docker exec -t $CONTAINERID python3 pre.py

# Segmentation with u-net
docker exec -t $CONTAINERID nnUNetv2_predict -i /app/input_pre/$SAMPLE_NAME -o /output -d 002 -c 2d -f all -device cpu

# Post-processing. It upsamples the masks from 512*512 to 2048*2048 and save an overlay version to check the segmentation quality. 
# If the original image size is not 2048*2048, you could add ``--size {YOUR_SIZE}` to resize it to the original size you wish.   
docker exec -t $CONTAINERID python3 post.py --alpha 0.20

# Copy the two segmentation to the output folder
docker cp $CONTAINERID:/output_real_value/ $ABS_PATH

# Stop and remove the docker container
docker stop $CONTAINERID
docker rm $CONTAINERID
```
