## Supplementary material for "A new protocol for multispecies bacterial infections in zebrafish and their monitoring through automated image analysis": S4 File Image analysis scripts: sort_images_lif.py

```
"""
LIF to TIFF Converter Script

This script is designed to convert Leica Image Files into individual TIFF files.
It processes each series, timepoint, channel, and Z-stack within the ND2 file and saves them as TIFF images.
The output is organized into directories for each series, with separate subdirectories for 
bright-field (C = 0) and fluorescence images (C > 0).

Sample1
	└── Images 
	│		├── Sample1_T00_C00_Z00.tif
	│		├── Sample1_T00_C00_Z01.tif
	│		└── ...
	│
	└── Fluor
			├── Sample001_T00_C01_Z00.tif
			├── Sample001_T00_C02_Z00.tif
			└── ...

Usage:
	python nd2_to_tiff_converter.py [path_to_nd2_file]
"""


import os
import argparse
import logging
from readlif.reader import LifFile

def create_directory(path):
	"""
	Create directory if it doesn't exist.
	"""
	if not os.path.isdir(path):
		os.makedirs(path)

def process_images(file_path):
	"""
	Process and save images from ND2 file.
	"""
	if not file_path.endswith('.lif'):
		logging.error("The provided file is not a LIF file.")
		return

	if not os.path.exists(file_path):
		logging.error("The provided file path does not exist.")
		return

	out_dir , _= os.path.splitext(file_path)

	try:
		lif = LifFile(file_path)
	except Exception as e:
		logging.error(f"Error reading LIF file: {e}")
		return


	for img in lif.get_iter_image():

		sample_name = img.name.split('/')[-1]
		
		fluor_dir = os.path.join(out_dir, sample_name, 'Fluor')
		img_dir = os.path.join(out_dir, sample_name, 'Images')

		create_directory(fluor_dir)
		create_directory(img_dir)

		for t in range(img.dims.t):
			for c in range(img.channels):
				for z in range(img.dims.z):
					img_name = f"{sample_name}_T{t:02}_C{c:02}_Z{z:02}.tif"
					frame = img.get_frame(z=z, t=t, c=c)
					if c == 0:
						frame.save(os.path.join(img_dir, img_name))
					else:
						frame.save(os.path.join(fluor_dir, img_name))
		logging.info(f"Saved {sample_name} to {os.path.join(out_dir, sample_name)}")


def main():
	"""
	Run the script with command line arguments.
	"""
	logging.basicConfig(level=logging.INFO)
	parser = argparse.ArgumentParser(description="Convert LIF file into individual tiff files")
	parser.add_argument("file_path", help="Path to LIF file")
	args = parser.parse_args()
	process_images(args.file_path)
	
if __name__ == "__main__":
	main()
```
